# Linking Aβ and tau in the amyloid cascade through the intersection of their proteostasis networks

**DOI:** 10.1101/2025.03.28.646065

**Authors:** Christine M Lim, John Hardy, Michele Vendruscolo

## Abstract

Amyloid plaques and neurofibrillary tangles are molecular hallmarks of Alzheimer’s disease. According to the amyloid cascade hypothesis, aberrant Aβ and tau behaviours contribute synergistically to accelerate the Alzheimer’s pathology. However, the complex molecular mechanisms linking Aβ and tau dysregulation remain to be fully characterised. To address this problem, we investigated the connection between Aβ and tau through the protein homeostasis (proteostasis) network. We asked whether Aβ proteostasis is linked to tau proteostasis. To answer to this question, we first mapped the proteostasis networks of Aβ and tau, and then studied the interplay of these two networks, identifying the molecular chaperone HSP90 as a central hub. To test this hub role of HSP90, we observed in a cell model that HSP90 and its co-chaperone SUGT1 mediate tau phosphorylation via GSK-3β in an Aβ42-dependent manner. Furthermore, we also observed that in turn HSP90 and SUGT1 increase the intracellular concentration of Aβ42. These results suggest that the HSP90/SUGT1 system may act as a hub in the amyloid cascade by lying at the intersection of the Aβ and tau proteostasis networks.

## Introduction

Alzheimer’s disease (AD) is a neurodegenerative disorder characterized by progressive cognitive decline and memory loss (1, 2). At the histopathological level, two hallmarks of AD are the accumulation of Aβ into amyloid plaques and of hyperphosphorylated tau into neurofibrillary tangles (3, 4). According to the amyloid cascade hypothesis, abnormal accumulation and aggregation of Aβ serve as critical initiating events, triggering a series of downstream pathological processes—including tau post-translational modifications, synaptic dysfunction, neuroinflammation, and ultimately neuronal death and cognitive decline (5-9). This hypothesis has gained renewed attention with recent clinical developments, notably the FDA approval of three antibodies targeting amyloid plaques (10-16).

Despite this clinical progress, the precise molecular mechanisms linking aberrant Aβ accumulation to tau dysregulation remain under investigation (9, 17-20). Various mechanisms have been proposed, including activation of tau-phosphorylating kinases (e.g. GSK-3β, CDK5, MAPK1/ERK2 and MAPK3/ERK1) (21-37), calcium signaling dysregulation, mitochondrial dysfunction, reactive oxygen species (ROS) production, and neuroinflammation (5-9). However, the complex interplay between these mechanisms has made it challenging to determine how they integrate into a coherent pathological cascade.

To bridge this critical gap, we asked whether within the amyloid cascade Aβ and tau could be linked through the protein homeostasis (proteostasis) system, which is ensemble of cellular processes that regulates the behaviour of proteins in terms of their conformations, interactions, concentrations and localizations (38, 39). This approach was prompted by the observation that the dysregulation of the proteostasis system has been associated with a wide range of neurodegenerative disorders (38-40). In this view, perturbations in the Aβ proteostasis network may affect the tau proteostasis network. To test this possibility, we first identified the proteostasis networks of Aβ and tau and then analysed their links.

By adopting this approach, we identified the molecular chaperone HSP90 and its co-chaperone SUGT1 as central players at the intersection of the proteostasis networks of Aβ and tau **(Figure 1)**. In the shared proteostasis network of Aβ and tau, HSP90 and SUGT1 lie on the shortest path between Aβ and tau, which includes also GSK-3β. The involvement of HSP90 has long been investigated in AD, since this molecular chaperone is a master regulator of a range of essential cellular processes such as survival, heat shock, protein folding, and client degradation (41, 42). Our analysis suggests that Aβ may alter the HSP90/SUGT1 system and promote its ability to promote the activity of GSK-3β in phosphorylating tau **(Figure 1)**. We tested this prediction in a cell model, finding that HSP90 and SUGT1 levels were perturbed in the presence of Aβ42 and that an Aβ42-dependent increase in phosphorylated tau (p-tau) required both HSP90 and SUGT1. By clarifying these proteostasis interactions, our findings help connect different molecular mechanisms underlying AD pathology, potentially opening new avenues for targeted therapeutic interventions.

**Figure 1.**
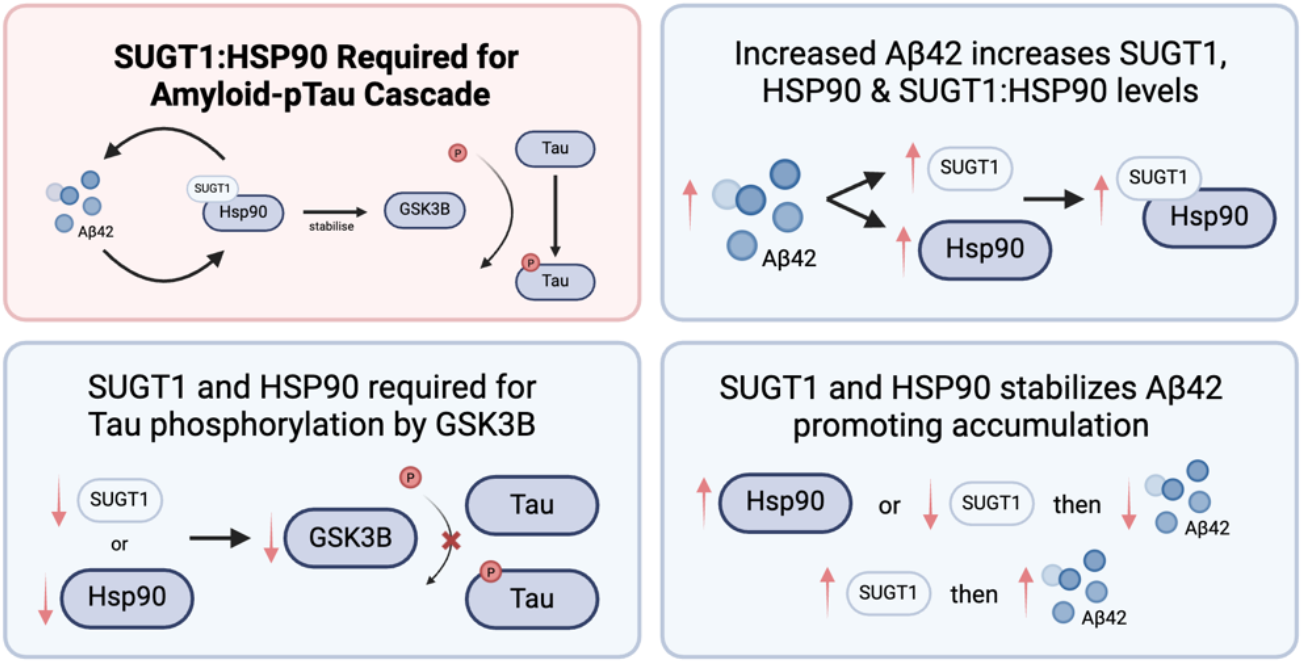
Schematic illustration of the modulation of the Aβ42/tau cascade by HSP90 and its co-chaperone SUGT1. We observed an increase in the SUGT1:HSP90 association in the presence of Aβ42, which in turn was linked with an increase in the tau-phosphorylating kinase GSK-3β. Upon treatment with exogenous Aβ42 or Aftin-5, we found an increase in SUGT1, HSP90, GSK-3β and p-tau levels **(Figure S1)**. The increased levels of Aβ42 were correlated with increased SUGT1:HSP90 association. This interaction appeared to be required for GSK-3β-induced tau phosphorylation as knockdown of either *SUGT1* or *HSP90* resulted in decreased GSK-3β and p-tau levels. Furthermore, increased levels of SUGT1 and HSP90 promoted the accumulation of Aβ42.

## Results

### The proteostasis networks of tau and Aβ

To study how Aβ affects tau via the proteostasis system, we identified the tau proteostasis network **(Figure 2A)** and the Aβ proteostasis network **(Figure 2B)**. For this purpose, we first obtained the proteostasis network components (43, 44) from the Proteostasis Consortium (https://www.proteostasisconsortium.com/). We then used the functional pairwise interactions from the Reactome database (45) (https://reactome.org/) to select all the interactions involving a proteostasis protein and Aβ or tau. We next filtered for inward interactions and bi-directional interactions, obtaining the final list of proteostasis proteins in the proteostasis network of Aβ (**Supplementary Table 1**) and in the proteostasis network of tau (**Supplementary Table 1**).

**Figure 2.**
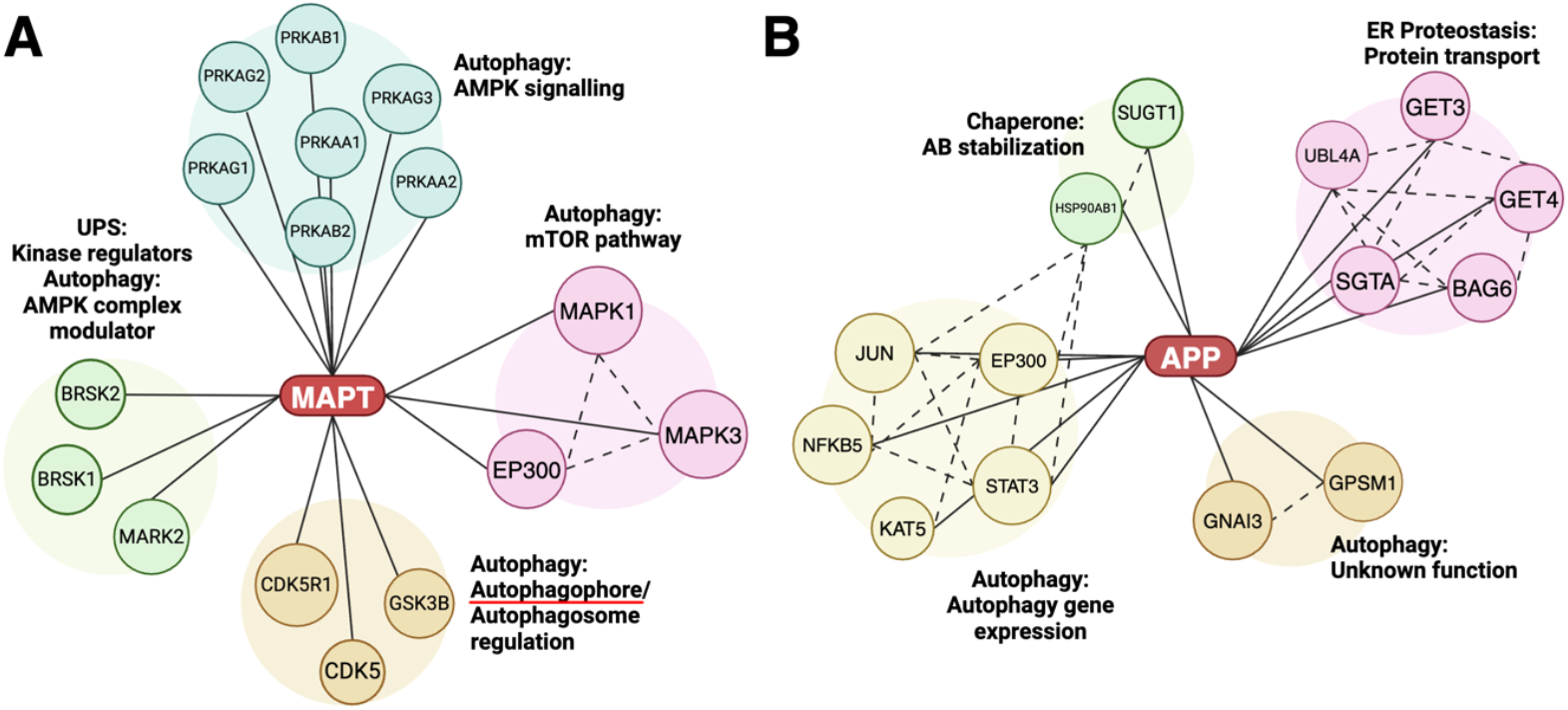
Proteostasis networks of Aβ and tau. **(A,B)** To study the mechanism by which Aβ promotes tau hyperphosphorylation by perturbing tau homeostasis, we identified the proteostasis network of tau (*MAPT*) (A) and the proteostasis network of Aβ (*APP*) (B). This analysis is consistent with an important role of autophagy in maintaining tau homeostasis, and with a contribution of molecular chaperones and protein transport to stabilising and transporting Aβ in addition to clearance via autophagy.

The proteostasis network of tau, a protein encoded by the *MAPT* gene, comprises 16 proteins, which are involved in the ubiquitin-proteosome system through kinase regulators (MARK2, BRSK1 and BRSK2) and in autophagy through four different processes: AMPK signalling (PRKAA1, PRKAA2, PRKAB1, PRKAB2, PRKAG1, PRKAG2 and PRKAG2), AMPK complex regulation (MARK2, BRSK1 and BRSK2), mTOR pathway (MAPK1, MAPK2 and EP300), and autophagophore/autophagosome regulation (CDK5, CDK5R1 and GSK-3β).

The proteostasis network of Aβ, a peptide derived from the proteolytic cleavage of amyloid precursor protein (APP), a protein encoded by the *APP* gene, comprises 14 proteins, which are involved in autophagy through gene expression (EP300, JUN, KAT5, NFKB1 and STAT3) and other processes (GNAI3 and GPSM1), in endoplasmic reticulum (ER) proteostasis through protein transport (BAG6, GET3, GET4, UBL4A and SGTA) and in molecular chaperones through Aβ stabilization (HSP90AB1 and SUGT1).

### Tau-phosphorylating kinases link the proteostasis networks of Aβ and tau

To identify connections between Aβ and tau through their proteostasis networks, we looked for all shortest paths from Aβ to tau through interactions between the proteins in their proteostasis networks. We found that the main connectors between Aβ and tau are the tau-phosphorylating kinases CDK5, GSK-3β, MAPK1 and MAPK3.

Since it was shown that Aβ activates GSK-3β (25, 46), which in turn phosphorylates tau, we further refined our focus on finding mechanistic links between Aβ and GSK-3β. The shortest paths from Aβ to tau via GSK-3β are visualized in **Figure 3**. This analysis revealed that HSP90 and its co-chaperone SUGT1 lie on the shortest path between Aβ and GSK-3β **(Figure 3)**. Since HSP90 is a member of both Aβ and tau proteostasis networks, we investigated whether increased Aβ levels result in hyperphosphorylation via altering tau proteostasis by studying the Aβ-HSP90:SUGT1-GSK-3β system. Three other Aβ interactors are found on the shortest path between Aβ and GSK-3β: MAPK14, RELA, and NFKB1 **(Figure 3)**.

**Figure 3.**
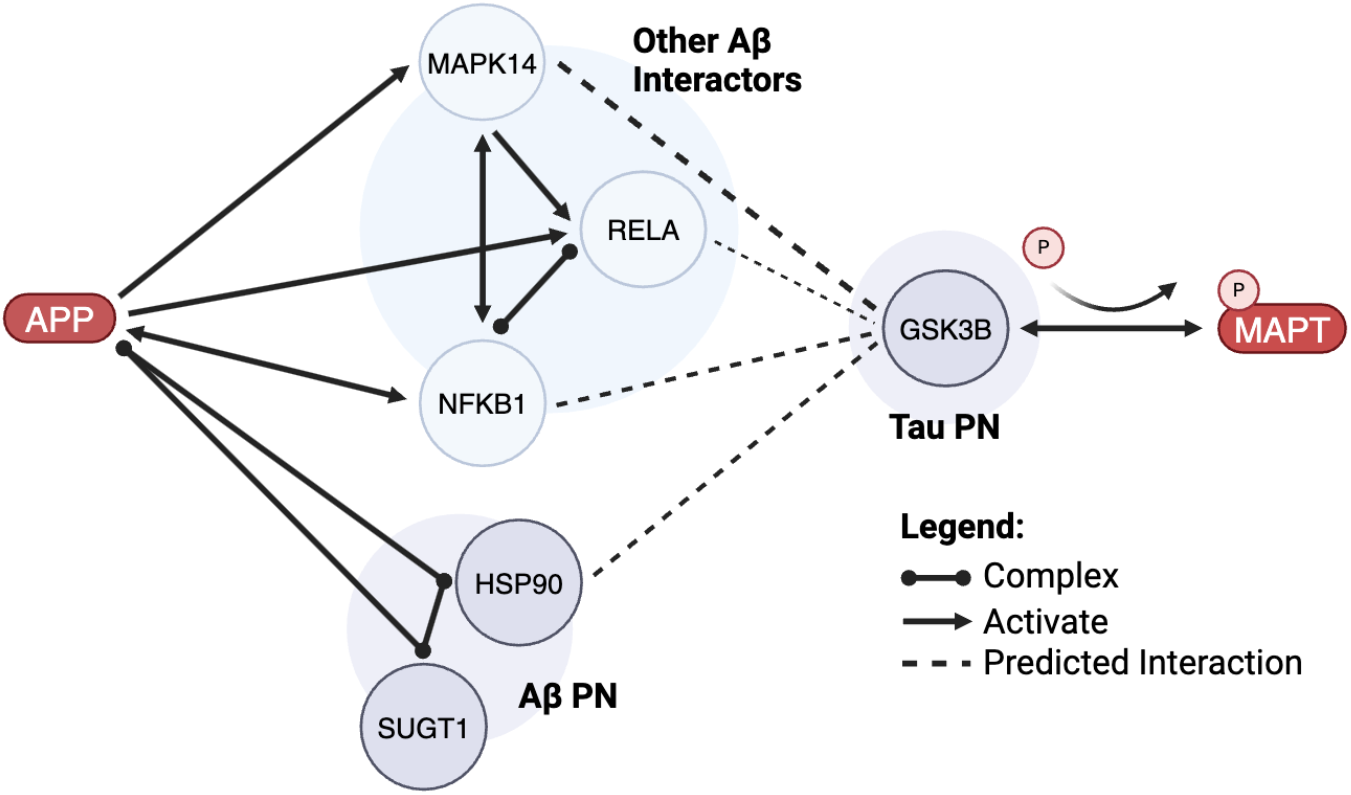
Tau-phosphorylating kinases link the proteostasis networks of Aβ and tau. To propose a possible mechanism for the cascade of Aβ to tau phosphorylation, we mapped the shortest paths between Aβ and tau **(Supplementary Table 1)**. The interactions are distinguished between complex formation (Complex), functional activation (Activate) and predicted (Predicted). The shortest path between Aβ and tau goes through GSK-3β, a kinase that phosphorylates tau. Five key proteins lie along the shortest path between Aβ and GSK-3β: the transcription factors MAPK14, RELA and NFKB1, and the molecular chaperones HSP90 and SUGT1.

### HSP90 modulates Aβ42-dependent tau phosphorylation via GSK-3β in SH-SY5Y cells

We found that increased phosphorylation of tau is associated with increased Aβ42 production in SH-SY5Y cells. When treated with Aftin-5, a compound known to increase the levels of Aβ42 (47), SH-SY5Y cells exhibited an increase in Aβ42 (**Figure S1**). This increase in Aβ42 was accompanied by elevated levels of p-tau (**Figure S1**). A similar increase in levels of p-tau was observed upon treatment of SH-SY5Y with exogenous monomeric Aβ42 (**Figure S2**). We then found that HSP90 inhibition prevented Aβ42-dependent tau phosphorylation. Given that HSP90 levels increased with increased Aβ42 levels (**Figure S1**) and that Aβ42 levels were correlated with HSP90 levels in SH-SY5Y cells treated with Aftin-5 (**Figure S1**), we hypothesized that Aβ42-dependent tau phosphorylation could be associated with HSP90. This hypothesis was further supported by colocalization and correlation of p-tau and HSP90 in Aftin-5-treated cells (**Figure 4A,B**). We tested this hypothesis via the addition of HSP90 inhibitor geldanamycin (GA) into SH-SY5Y cells treated with Aftin-5 which prevented the increase in tau phosphorylation (**Figure 4A,C,D**), possibly because HSP90 inhibition lowers levels of GSK-3β in SH-SY5Y cells treated with Aftin-5 **(Figure 4E,F)**.

**Figure 4.**
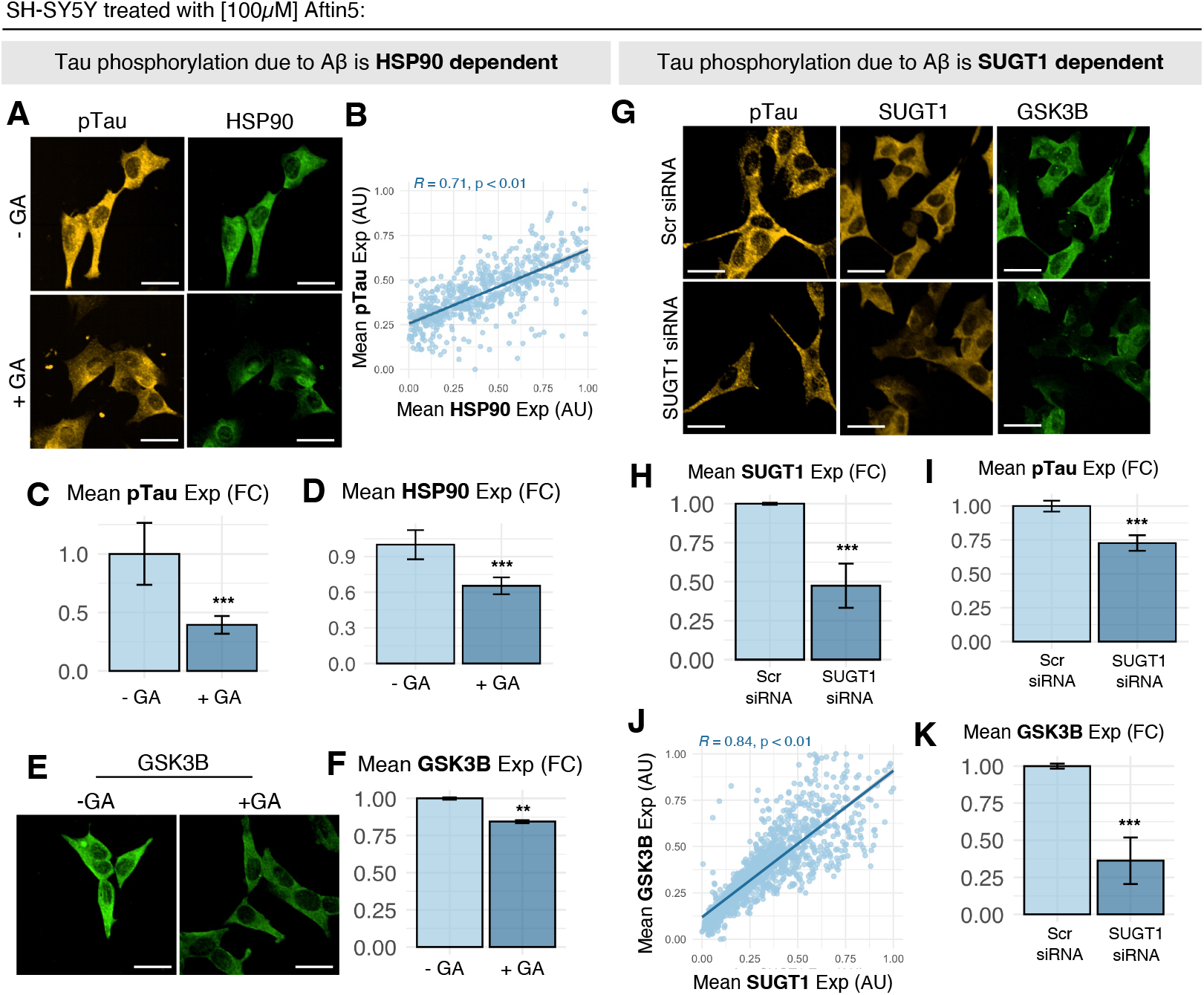
Increased Aβ42 levels correlate with increased tau phosphorylation and are associated with increased HSP90 and SUGT1 levels. **(A)** Representative images of SH-SY5Y cells treated with Aftin-5, which increases Aβ42 levels and GA, which inhibits HSP90. **(B-D)** Single-cell quantification revealed that p-tau levels are correlated with HSP90 levels (B), and that an increase in tau phosphorylation **(Figure S1)** can be prevented by HSP90 inhibition using GA at 10 nM (C,D). **(E**,**F)** Inhibition of HSP90 also results in decreased levels of GSK-3β. All GA experiments were conducted with N=3 biological replicates of n=3 technical replicates. **(G)** Representative images of Aftin-5-treated SH-SY5Y cells with SUGT1 KD. **(H-K)** With decreased levels of SUGT1 (H), the increase in p-tau **(Figure S1)** due to treatment with Aftin-5 was reduced (I). This was likely because GSK-3β levels are correlated with SUGT1 levels (J), and GSK-3β levels are lower in Aftin-5 (+) cells treated with SUGT1 siRNA compared to controls (K). All SUGT1 KD experiments were conducted with N=3 biological replicates of n=2 technical replicates.

### The HSP90 co-chaperone SUGT1 enhances HSP90-mediated GSK-3β activity

Since HSP90 is a master regulator of molecular chaperone activities involved in multiple pathways, we explored the role of the HSP90 co-chaperone SUGT1 in GSK-3β activity. To do this, we knocked down *SUGT1* in SH-SY5Y cells treated with Aftin-5 **(Figure 4G)** and found that lower levels of SUGT1 **(Figure 4H)** were associated with lower levels of p-tau **(Figure 4I)**, possibly because SUGT1 levels were in turn correlated with GSK-3β levels **(Figure 4J)**. Given that: (i) SUGT1 colocalised with HSP90, (ii) SUGT1 knockdown were correlated with lowered levels of HSP90:SUGT1, and (iii) SUGT1 knockdown mimicked the effect HSP90 inhibition on modulating GSK-3β levels, our results suggest that SUGT1 promotes an HSP90-dependent increase in GSK-3β levels.

### HSP90 and SUGT1 increase Aβ42 levels

The findings described above indicate that under perturbing conditions, the HSP90 and SUGT1 promote tau hyperphosphorylation. We hence studied the role of HSP90 and SUGT1 in Aβ42 regulation in control cells. Cells treated with increasing concentrations of GA, an HSP90 inhibitor (**Figure 5A**) showed lower levels of intracellular Aβ42 (**Figure 5B**), which correlated with lower levels of HSP90 (**Figure 5C**). We hence inferred that under normal conditions, HSP90 promoted the maintenance of intracellular Aβ42. Similarly, cells transfected with SUGT1 cDNA (**Figure 5D**) expressed higher levels of SUGT1 (**Figure 5E**) and had higher levels of intracellular of Aβ42 (**Figure 5F**). These results suggest that under normal conditions, SUGT1 is also required for the maintenance of intracellular Aβ42.

**Figure 5.**
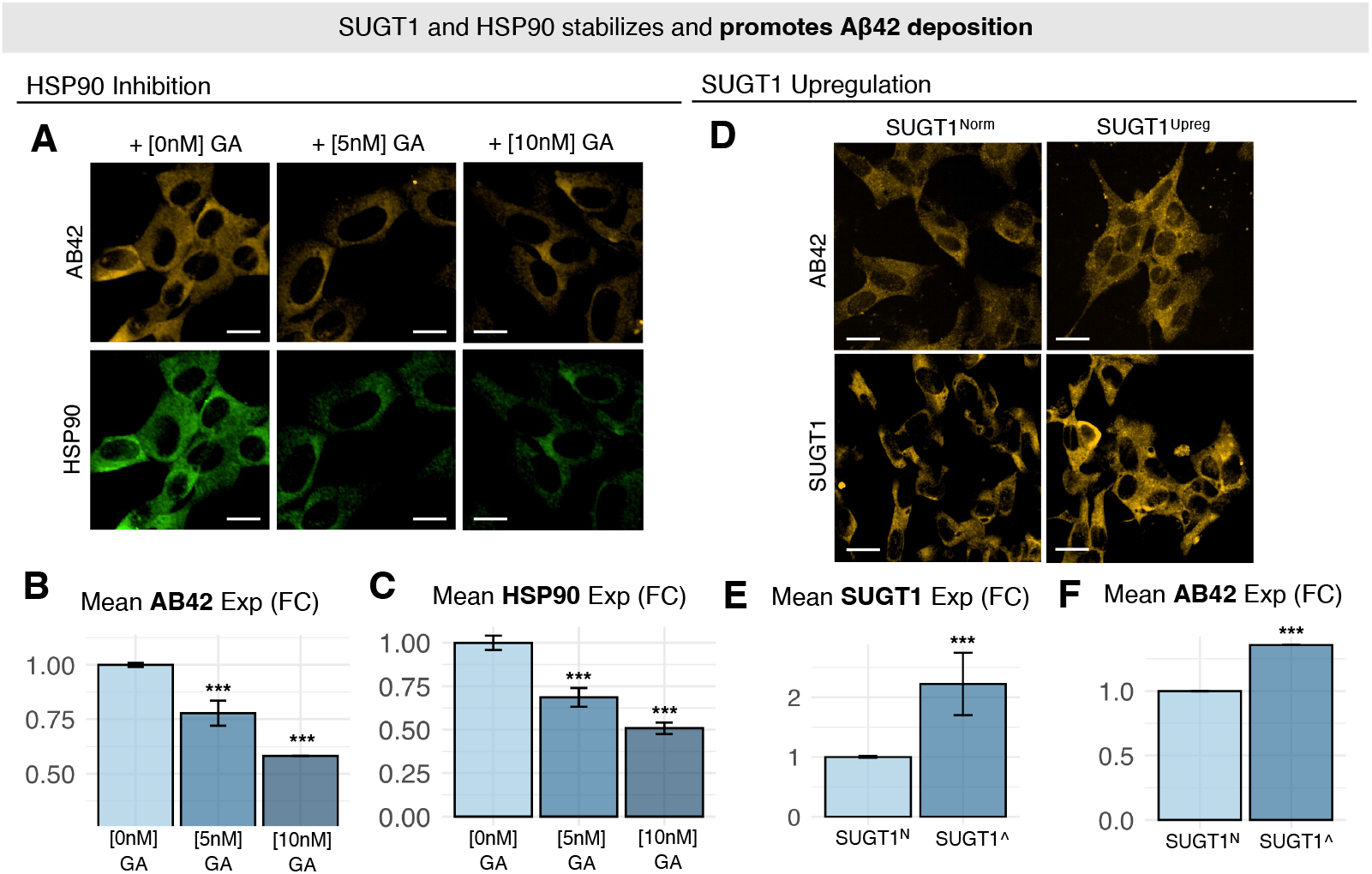
HSP90 and SUGT1 increase the intracellular concentration of Aβ42. **(A)** Representative images of SH-SY5Y cells treated with increasing concentrations of geldanamycin (GA), an HSP90 inhibitor. **(B**,**C)** Intracellular levels of Aβ42 decrease (B) with increasing HSP90 inhibition (C). **(D)** Representative images of SH-SY5Y cells with SUGT1 upregulation via transfection are shown. **(E**,**F)** Increased levels of SUGT1 (E) resulted in increased intracellular levels of Aβ42 (F). N=3 biological replicates of n=2 technical replicates.

### Knockdown of the HSP90 co-chaperone FKBP51/*FKBP5* increases GSK-3β levels

To check if other HSP90 co-chaperones also had the ability of reduce GSK-3β, we looked for alternative upregulated HSP90 co-chaperones. Our comparison of 4 microarray studies revealed that *FKBP5* is consistently upregulated in AD patients **(Figure S3A)**. We corroborated this information with literature and found that *FKBP5* expression was found to increase with aging and elevated *FKBP5* levels have also been reported in AD patients (48, 49). Furthermore, FKBP51 levels have also been reported to correlated with accumulation of tau oligomers (50). Given these observations, we first confirmed that FKBP51 was upregulated at the protein level in response to Aβ42 using our assay where SH-SY5Y cells were treated with Aβ42 **(Figure S3B**,**C)**. We then knocked down *FKBP5* via treatment with *FKBP5* siRNA **(Figure S3D)**. We found that *FKBP5* knockdown resulted in resulted in a further increase in GSK-3β levels upon Aβ42 treatment **(Figure S3E**,**F)**.

### SUGT1 levels are higher in AD-vulnerable brain regions

The results described above suggest that SUGT1 elevation promotes the stabilization by HSP90 of GSK-3β, thus enhancing tau phosphorylation. To study this effect, we compared SUGT1 levels in the entorhinal cortex (aa AD-vulnerable brain region) vs the cerebellum (a non-AD-vulnerable brain region) in normal humans, mice, and pigs. Our comparison reveals that AD-vulnerable brain regions have a higher SUGT1 level across all 3 organisms compared to less AD-vulnerable regions **(Figure S4)**.

## Discussion

In this work, we investigated whether Aβ and tau could be linked within the amyloid cascade through the proteostasis system. By mapping the proteostasis networks of Aβ and tau, we identified the HSP90:SUGT1 as a critical hub lying at their intersection. Our experiments indicated that Aβ42 elevated the levels of both HSP90 and SUGT1 in SH-SY5Y cells, and were associated with an enhanced activity of the kinase GSK-3β, which promotes tau hyperphosphorylation. We also found a reciprocal effect, wherein the HSP90:SUGT1 complex further increases intracellular levels of Aβ42. These observations suggest a self-reinforcing feedback loop that potentially accelerates AD pathology.

These findings complement a body of literature exploring molecular chaperones, particularly HSP90, as therapeutic targets in AD due to their critical roles in protein folding and clearance of misfolded aggregates. (51). Previous studies have demonstrated the efficacy of both N-terminal (e.g., geldanamycin, 17-AAG) and C-terminal (e.g., novobiocin derivatives) HSP90 inhibitors in reducing Aβ toxicity and tau accumulation (41, 51-57). We note, however, that the global inhibition of HSP90 remains challenging because of its extensive involvement in various essential cellular processes, which can lead to off-target effects. Our work provides an alternative by highlighting the potential of modulating the specific HSP90-co-chaperone interactions. In particular, the HSP90 co-chaperone SUGT1 could serve as a more selective therapeutic target. However, despite indicating that SUGT1 promotes GSK-3β-induced tau phosphorylation, our current study does not fully establish the specificity of the SUGT1– HSP90 interaction. Future experiments, such as structural characterization, direct binding assays, and mutational analyses, will be essential to validate SUGT1 as a viable therapeutic target and to demonstrate specificity.

Additionally, our discovery of a bidirectional interaction between HSP90:SUGT1 and Aβ42 levels suggests that disrupting this feedback loop could be particularly beneficial in slowing or halting disease progression. This observation underscores the potential advantage of selectively targeting intersection nodes within proteostasis networks, rather than broadly inhibiting proteostasis components.

The possible roles of other co-chaperones, such as FKBP51, also emerged in our investigation. Although our results showed increased GSK-3β levels following *FKBP5* knockdown, the precise functional role of FKBP51 within the proteostasis system and its relationship to the HSP90:SUGT1 complex remains unclear. Further clarification of the interaction of FKBP51 with HSP90 and its impact on tau and Aβ42 dynamics may illuminate additional therapeutic strategies or help refine existing ones.

In conclusion, our study identifies the HSP90:SUGT1 complex as a key intersection between Aβ and tau proteostasis networks, providing a mechanistic understanding of how Aβ accumulation drives tau hyperphosphorylation through proteostasis dysregulation. The discovery of a reciprocal feedback loop, where HSP90:SUGT1 simultaneously exacerbates tau pathology and increases intracellular Aβ42, underscores the therapeutic value of selectively disrupting this pathological circuit. Future research should prioritize characterizing the specificity of HSP90–SUGT1 interactions and explore co-chaperone modulation strategies, potentially overcoming the limitations of global proteostasis inhibitors currently under investigation.

## Materials and Methods

### Proteostasis network components

The proteostasis network components (43, 44) were obtained from the Proteostasis Consortium (https://www.proteostasisconsortium.com/).

### Protein-protein interaction (PPI) network

Functional pairwise interactions (version 2021) were downloaded from the Reactome database (45) (https://reactome.org/).

### Proteostasis network of tau

To identify the proteostasis network of tau, we filtered the pairwise interactions for any interactions involving a proteostasis protein and tau (*MAPT*). We then filtered those interactions for bi-directional interactions such as complex formations or inward interactions. The remaining proteostasis interactors for each of tau (*MAPT*) form the final list of proteostasis proteins involved in tau proteostasis **(Supplementary Table 1)**. A literature search was carried out to annotate and classify the proteostasis proteins for their regulatory activity on tau (**Supplementary Table 1**). All other proteins (not included within our list of proteostasis network components) exerting inward functional interactions on tau are also separately available in **Supplementary Table 1**.

### Proteostasis network of Aβ

To identify the proteostasis network of Aβ (*APP*), we carried out the same procedure described above for tau (**Supplementary Table 1**). All other proteins (not included within our list of proteostasis network components) exerting inward functional interactions on Aβ are also separately available in **Supplementary Table 1**.

### Cell culture experiments

Human neuroblastoma cells (SH-SY5Y) were cultured in DMEM/F-12 GlutaMAX™ supplemented with 10% heat-inactivated fetal bovine serum (hiFBS) and maintained at 37 °C, 5% CO_2_, 95% relative humidity. For each experiment, SH-SY5Y cells were plated in PerkinElmer Cell Carrier Ultra plates at a density of 7.5k-10k cells/well. The cells were incubated for 24 h to allow adequate cell attachment before siRNA feeding. The cells were transfected with siRNA for 24 h before 48 h of Aftin-5/exogenous Aβ42 treatment. When Aftin-5 was used for treatment, Aftin-5 was replenished into the media every 24 h. Cells treated with Aftin-5 and GA were treated with GA after an initial 24 h incubation with Aftin-5. All cells were fixed using 4% paraformaldehyde (PFA) dissolved in D-PBS (+/+) and stored at 4 °C for less than a week before staining.

### Treatment of SH-SY5Y cells with SiRNA

During siRNA feeding, full culture medium was replaced with fresh DMEM/F-12 GlutaMAX™ with 1% hiFBS and fed with a final concentration of 10 nM siRNA (mixture of Thermo Fisher silencer select SUGT1 siRNAs: #s21444 and #s21446). Lipofectamine RNAiMAX was used as the transfection reagent. The cells were incubated with siRNA for 24 h before any further treatment.

### Preparation of Aβ42 monomers for the exogenous treatment of SH-SY5Y cells

All monomers used for exogenous treatment were freshly purified right before treatment. The monomers were purified from potential oligomeric species using size exclusion chromatography (Superdex 75 10/300 GL column, GE Healthcare, Chicago, IL). Recombinant Aβ42 was expressed and purified as previously reported. Lyophilized samples of Aβ42 were dissolved in 6 M GuHCl and 20 mM sodium phosphate buffer, pH 8 and incubated on ice for 1 h. 20 mM NaP buffer, pH 8 was used for elution during monomer purification. During purification, the flow rate was kept at 0.7 mL/min. Protein elution was tracked with the use of a chromatogram, assessing both the elution volume and the absorbance levels at λ=280 nm. Only monomers eluted at the peak were collected and used for treatment. The concentration of Aβ42 was determined from the absorbance of the integrated peak area (ε = 1,490 M^−1^ cm^−1^ and MW = 4,514.1 g/mol).

### ICC staining and imaging

Cells were permeabilized with 0.1% Triton™ X-100 (#85111, Thermo Fisher) for 20-30 min at room temperature then rinsed three times with D-PBS (+/+). The cells were then incubated for in 1.5% BSA blocking buffer for 1 h at room temperature. Following which, the cells were incubated with the appropriate dilution of the desired primary antibodies overnight at 4 °C. Primary antibodies used are as follows: Aβ42 (#44-344, Invitrogen), HSP90 (#ab13492, Abcam), GSK-3β (#MA5-15597, Invitrogen), p-tau S396 (#44-752G, Invitrogen), and SUGT1 (#PA5-28237, Invitrogen). Following overnight incubation with the relevant primary antibodies, the cells were washed three times with D-PBS (+/+) and incubated with secondary antibody Alexa Fluor 555 anti-rabbit (#A21428, Thermo Fisher) diluted in blocking solution for 1 h at room temperature. Finally, the cells were rinsed three more times with D-PBS (+/+) and stained with Hoechst 33342 (#H3570, Thermo Fisher) before imaging. All 96-well plates generated were imaged using the Opera Phenix High-Content Confocal microscope at a 40X magnification. Images of 24 or more unique frames per well for each of 3 biological replicates were acquired for quantification.

### Single-cell protein quantification

Image analysis and quantifications were performed with the Harmony High-Content Imaging and Analysis Software (Perkin Elmer). Image processing steps were as follows: (1) cell segmentation, (2) summation of raw protein fluorescence (for each channel) of all pixels in each cell mask, (3) quantification of total cell area for each cell mask, and (4) normalization of protein fluorescence as described below. All segmentation, fluorescence quantification, and cell area summation were done using the software default settings. To normalize the protein fluorescence and get average protein levels: average protein level = total protein fluorescence in a cell / total cell area of the cell. Single-cell outliers were determined and removed if their average protein levels were larger than 3Q+1.5x IQR from 3Q or smaller than 1Q-1.5x IQR. All fold changes were calculated against the relevant controls treated, stained and imaged at the same time within the same 96-well plate.

### Differential analysis of HSP90 co-chaperones in four microarray studies of AD patients

Four microarray studies were selected for differential expression (DE) analysis: GSE48350, GSE36980, GSE5281, and GSE4757. Microarray datasets available in the pandaomics database (https://pandaomics.com/access) that contained more than five control and five AD patient samples, sequenced for more than 12,000 genes were selected and included in the studies. Where samples specifically from the entorhinal cortex (EC) were available, only AD EC samples were compared against control EC samples. DE analysis was carried out using Limma. Log_2_ FC values are available in **Supplementary Table 2**.

### Comparison of SUGT1 levels in AD-vulnerable brain regions

SUGT1 normalised transcripts per million (nTPM) values were obtained from GTEx and the human protein atlas (HPA) dataset for pigs and mice. Two brain regions were compared: the entorhinal cortex/hippocampus for which AD is understood to originate in the brain (most AD-vulnerable), and the cerebellum that is last affected in AD progression (least AD-vulnerable). nTPM values used for this comparison are available in **Supplementary Table 3**.

## Supplementary Information

**Figure S1.**
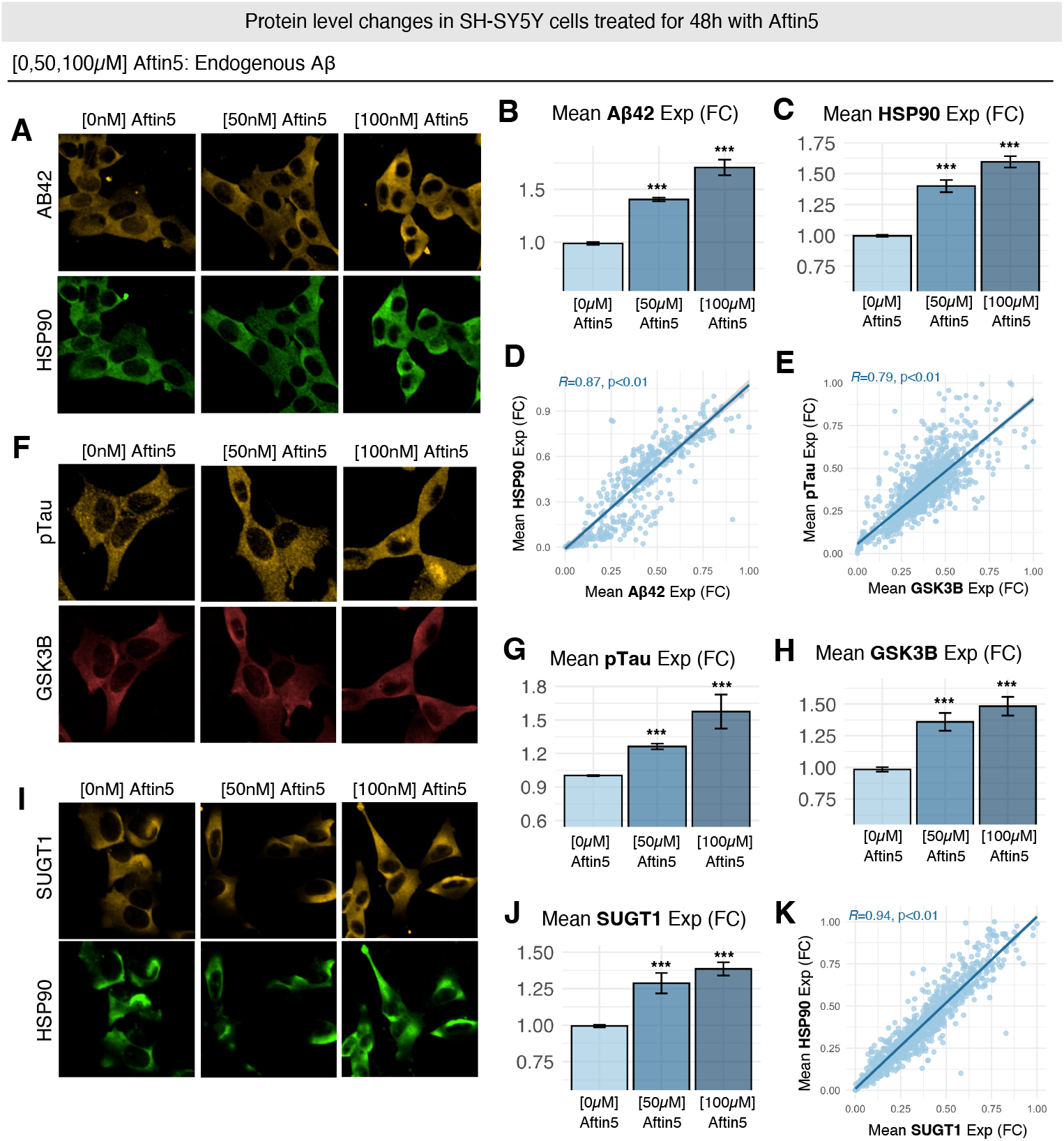
HSP90, p-tau, GSK-3β and SUGT1 levels increased with increasing levels of Aβ42 in SH-SY5Y cells treated with Aftin-5. **(A-C)** Upon treatment with increasing concentrations of Aftin-5, Aβ42 and HSP90 levels increased and the two proteins colocalised. **(D)** Cellular levels of Aβ42 and HSP90 were correlated suggesting that increased Aβ42 levels induced by the Aftin-5 treatment promoted HSP90 expression. **(E)** Under the same conditions, GSK-3β levels were also correlated with p-tau levels. **(F-H)** p-tau and GSK-3β colocalised and their average levels increased with increasing concentrations of Aftin-5 and Aβ42. **(I-K)** SUGT1 levels increased with increasing concentrations of Aftin-5, and SUGT1 colocalized with HSP90. SUGT1 and HSP90 were correlated, and cells with higher levels of HSP90 tended to have higher levels of SUGT1. N=3 biological replicates of n=3 technical replicates.

**Figure S2.**
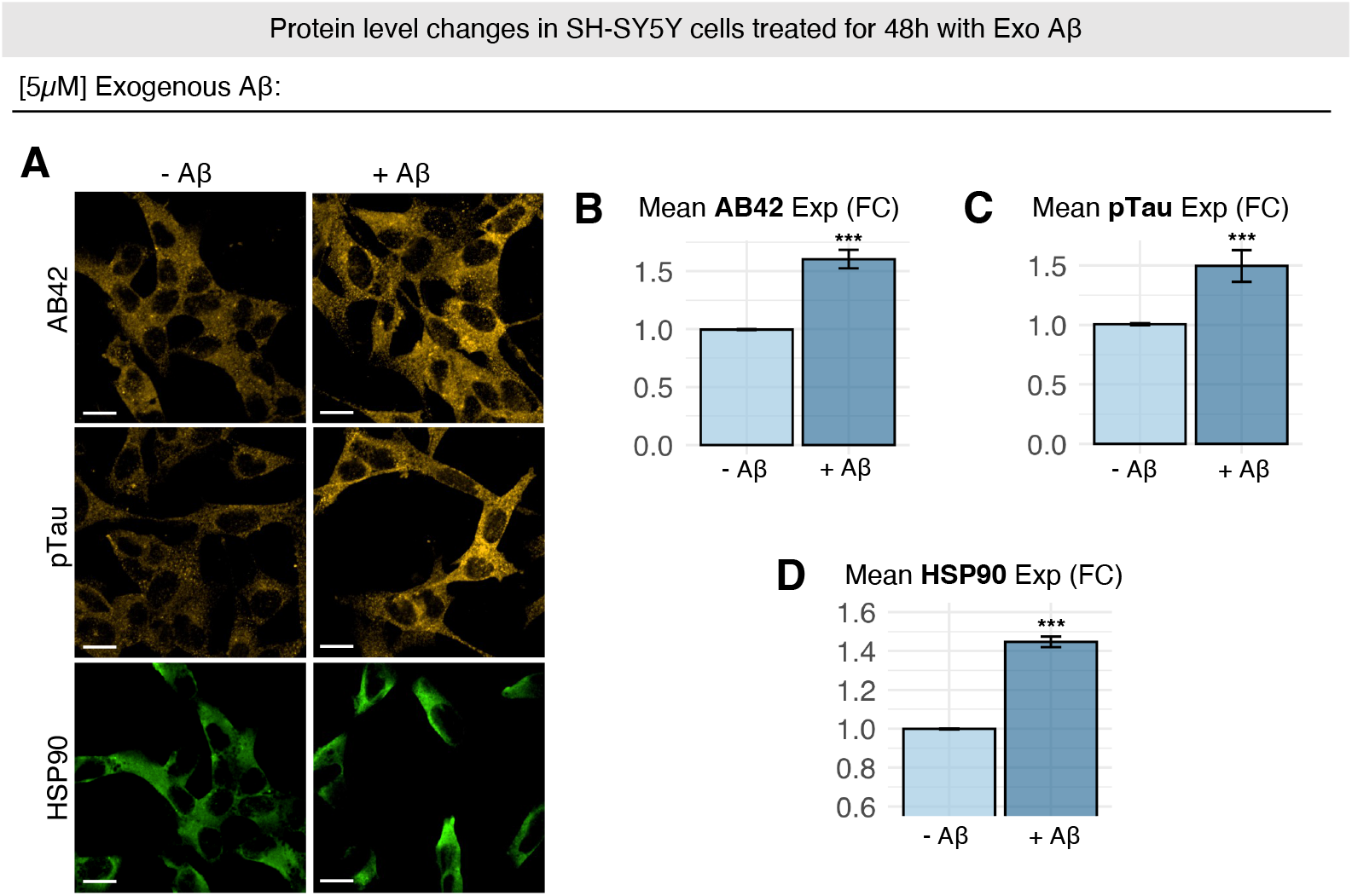
HSP90 and p-tau levels rose in SH-SY5Y cells treated with exogenous Aβ42. **(A)** Representative images of SH-SY5Y cells treated with exogenous Aβ42 at 5 μM concentration. **(B-D)** Treatment of SH-SY5Y cells with exogenous Aβ42 resulted in similar increases in Aβ42 (B), p-tau (C), and HSP90 (D). N=2 biological replicates of n=4 technical replicates.

**Figure S3.**
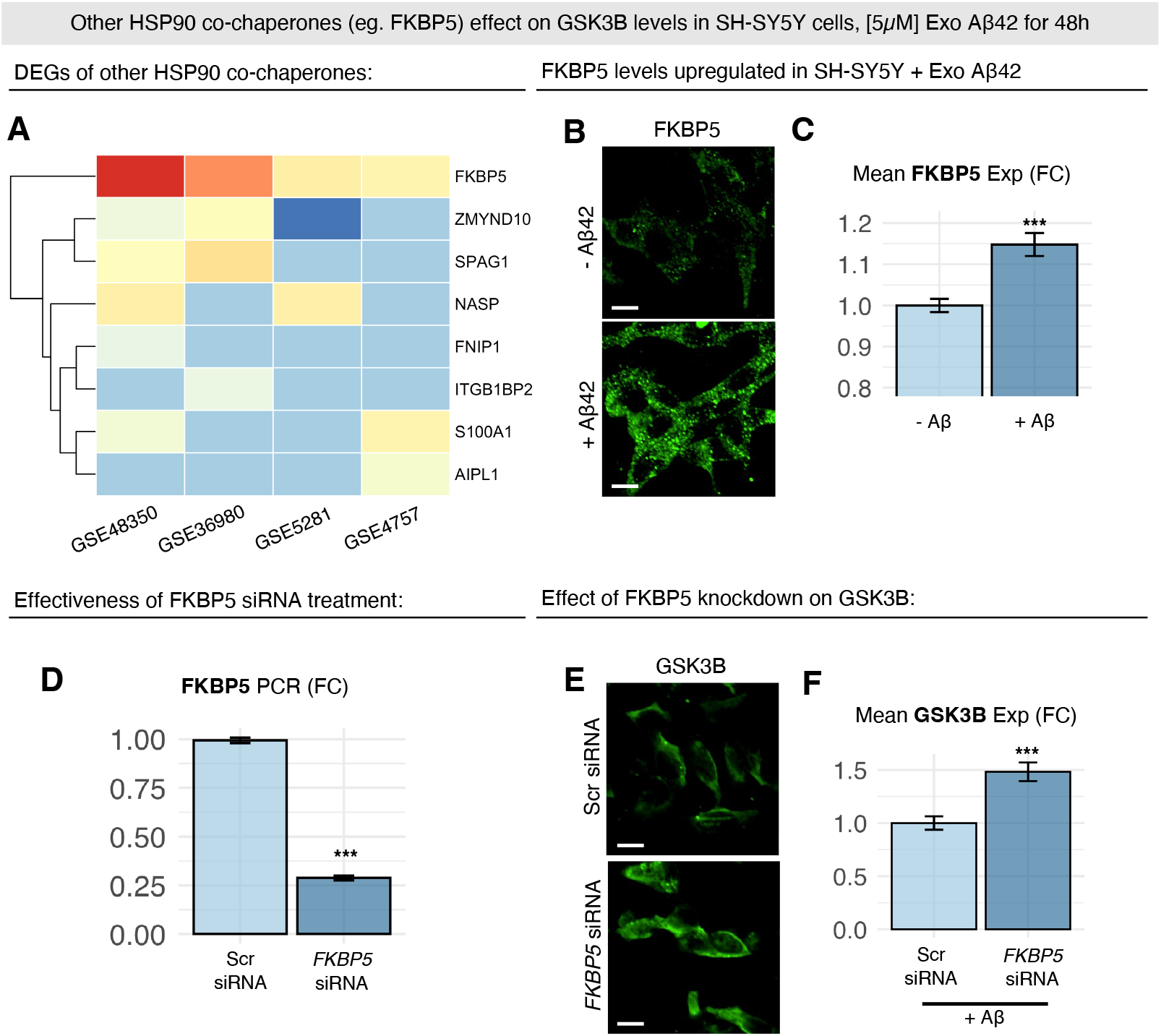
*FKBP5* knockdown increased GSK-3β levels. **(A)** Differential expression of HSP90 co-chaperones across four microarray datasets of AD patients. The heatmap presents differential expression log2 FC of HSP90 co-chaperones with log2 FC > 0.3 in at least one dataset. We found that the HSP90 co-chaperone FKBP51 was consistently upregulated in AD patients across all four datasets. **(B)** Representative images of FKBP51 levels in SH-SY5Y cells treated ± exogenous Aβ42 at 5 μM concentration. **(C)** Upon treatment with exogenous Aβ42, the mean FKBP51 expression increased. To measure the effects of FKBP51 on GSK-3β, we treated SH-SY5Y cells with *FKBP5* siRNA. **(D)** Validation of *FKBP5* knockdown via PCR. **(E)** Representative images of GSK-3β levels in SH-SY5Y cells treated with *FKBP5* siRNA and the scramble siRNA control are shown. **(F)** Knockdown of *FKBP5* increased GSK-3β levels. N=2 biological replicates of n=3 technical replicates.

**Figure S4.**
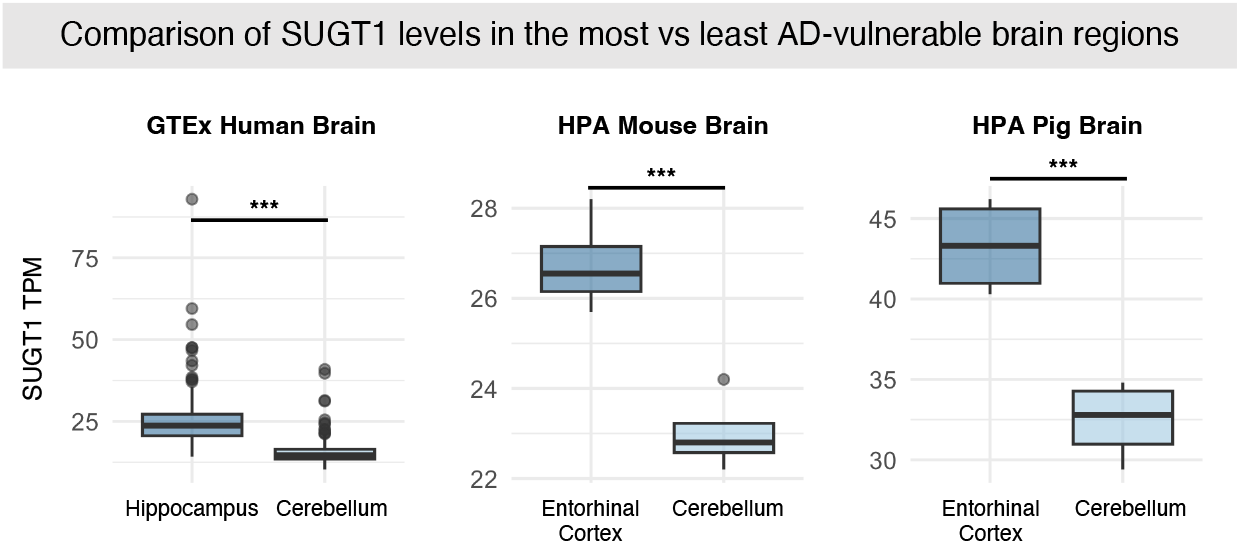
*SUGT1* levels are higher in AD-vulnerable brain regions but lower in AD-resistant brain regions. A comparison of early AD-affected regions (entorhinal cortex and hippocampus) against AD-resistant regions (cerebellum) revealed that AD-affected regions have higher levels of *SUGT1* across human, pig, and mouse.

